# High variability in the attractiveness of municipally-planted decorative plants to pollinators

**DOI:** 10.1101/2024.01.25.577170

**Authors:** Tomer J. Czaczkes, Carsten Breuss, Christoph Kurze

## Abstract

Insect populations are declining globally. A major driver of this decline is land use change, including urbanisation. However, urban environments can also offer a wide range of floral resources to pollinators, through ornamental plantings, but these can vary widely in their attractiveness to insects. Often, the largest single planter of ornamental plants in an urban area is the municipality. Here we evaluated the decorative plantings carried out by the city municipality of Regensburg, Germany, by systematically surveying insect visitations on different plant types in late summer, when forage is often limited for pollinators. We found a 130-fold increase from the least to the most attractive plants, and high variation in which insect groups were attracted to which plants. While honey bees, which are not conservation concern, were the most common insect visitors, some decorative plants attracted a very large proportion of wild bees, flies, and wasps. Our results demonstrate that there is great scope for increasing the supply of urban forage to pollinators in general, and specific groups in particular, without requiring new decorative plant types to be sourced or planted. We argue that providing local evidence-based guidance to municipalities offers a quick and potentially cost-neutral method for supporting urban insect populations.

## Introduction

Insect populations are declining globally (Goulson 2019; Wagner et al. 2021; Blüthgen et al. 2023). Of special interest is insect pollinator decline, as pollination is a key ecosystem service offered overwhelmingly by insects (Potts et al. 2010; Nath et al. 2023). Declining pollinator numbers and diversity is set to have large adverse effects on crop production, as well as threatening wild plant species which rely on insect pollination (Bauer and Wing 2010; Bauer and Sue Wing 2016; Pérez-Méndez et al. 2020). The reasons for insect pollinator decline are diverse (Potts et al. 2010), and include habitat loss due to land changes (Peters et al. 2022; Ganuza et al. 2022), pesticide impacts (Brittain et al. 2010), pathogen spillover from managed bees (Manley et al. 2015), and ecosystem changes caused by climate change (Vasiliev and Greenwood 2021), with habitat loss and degradation being perhaps the most important (Ganuza et al. 2022).

The loss and degradation of habitats, and subsequent impact on pollinators, is driven to a large extent by urban development and agricultural intensification (McKinney 2006; Winfree et al. 2009; Bates et al. 2011). This results, amongst other challenges, in a loss of floral resources (Fuller 1987; Potts et al. 2010; Kennedy et al. 2013; Baude et al. 2016; Jones et al. 2021). However, recent trends in land use changes may be having a positive effect on resource provision for pollinators; specifically, changes from intensive arable land use to low density urban landscapes. Urban landscapes can be a suitable habitat for pollinators since they often offer a high diversity of ornamental flowering plants (Baldock et al. 2015, 2019; Hall et al. 2017; Honchar and Gnatiuk 2020). Indeed, such ornamental flowering plants may be planted or bred to offer long periods of flower availability, well into seasons of low nectar flow (Salisbury et al. 2015). The important role of urban environments in supporting pollinators on heavily impacted landscapes is demonstrated by reports of a positive correlation between urbanisation and biodiversity, to a point (Lowenstein et al. 2014; Theodorou et al. 2017; Biella et al. 2022), although bumble bee colony health and performance are negatively correlated with percentage impervious surfaces (Theodorou et al. 2022). Ornamental plants may fill gaps in the nectar and pollen resources offered by wild plants (Salisbury et al. 2015; Seitz et al. 2020; Erickson et al. 2020). Indeed, cities often support a similar, or even larger, diversity of pollinators compared to agricultural landscapes (Baldock et al. 2015; Hall et al. 2017; Wenzel et al. 2020).

However, ornamental plants vary widely in their usefulness to pollinators. Some ornamental plant cultivars, such as double flowers (which have an extra set of petals instead of anthers) offer no nectar or pollen, or have inaccessible nectarines (Comba et al. 1999; Corbet et al. 2001; Garbuzov et al. 2015; Honchar and Gnatiuk 2020). There is a huge variation in attractiveness to pollinators between ornamental plants. For example, Garbuzov et. al (2015) report a 100-fold difference between the most and least attractive ornamental flowers they surveyed. Even amongst ornamental flowers chosen to be attractive to pollinators, between a 5- and 40-fold difference in attractiveness from most to least attractive was reported (Marquardt et al. 2021a, b; Palmersheim et al. 2022). Indeed, most ornamental plants sold in UK garden centres are not attractive to pollinators (Garbuzov et al. 2017). Additionally, different plants and cultivars are differentially attractive or useful to different insect groups (Rollings and Goulson 2019; Erickson et al. 2021), with many ornamental cultivars being especially popular with generalist insect species not in need of support, such as *Apis meliffera* and *Bombus terrestris* (Rollings and Goulson 2019; Honchar and Gnatiuk 2020; Peters et al. 2022). However, the reverse situation also exists – for example, *Anthemis tinctoria* was reported to attract a wide variety of pollinators such as wild bees, flies, and butterflies, but not honey bees or bumble bees (Rollings and Goulson 2019). While various floral traits have been found to correlate with attractiveness to pollinators (such as floral area cover (Grindeland et al. 2005; Marquardt et al. 2021a; Kalaman et al. 2022), colour (Reverté et al. 2016; Chapman et al. 2023), shape (Howard et al. 2019; Erickson et al. 2022; Chapman et al. 2023), and nutritional offering (Comba et al. 1999; Corbet et al. 2001; Chapman et al. 2023)), the huge variety of cultivars and interaction with local conditions make accurate prediction of cultivar attractiveness difficult. As such, Rollings and Goulson (2019) recommend that tests for the attractiveness of a wide variety of ornamental cultivars be carried out, ideally at various sites with varying environmental conditions.

Many municipalities manage extensive decorative flower beds. Such flower beds are often replanted multiple times a year, to provide constant blooms. The municipality is thus often the biggest planter of decorative plants in an area. Frequent replanting means that it can be agile in terms of optimising plantings, and it offers a single entity with which to interact. It is thus a highly attractive partner when attempting to provide evidence-based advice for improving food provision for urban pollinators. The first step in such an attempt is to gather local data on ornamental flower visitations by pollinators in municipally-planted flower beds. Thus, the aim of this study is to provide this data. We surveyed insect presence on broad variety of annual ornamental flowers planted by the City of Regensburg municipality, in order to both provide more quantitative data on which plants currently being deployed as ornamentals are most attractive to pollinators, and in order to provide the municipality data on which plant species they should favour in future planting cycles.

## Methods

The study was carried out over 11 public parks and public places in Regensburg, Germany (see table 1). Study locations ranged in from the centre to the edge of the city (2.5 km away from city centre). In total, 30 morphogroups (henceforth plants) were examined, including 25 perennials, 1 biennial, and 4 annuals. Four morphogroups contain two different cultivars or species, as they could not be identified before the start of the study (see supplement table TS1). Note that as we could not control what plants were available in each location, the distribution of different plants over the study areas is uneven. The study was carried out from 16.08.2023 to 06.09.2023. Observations were made between 8:00 – 11:00 in the morning and 14:00-17:00 in the afternoon, on days when it was not raining or excessively windy. Data were only collected from plants in full bloom.

**Table 1:**
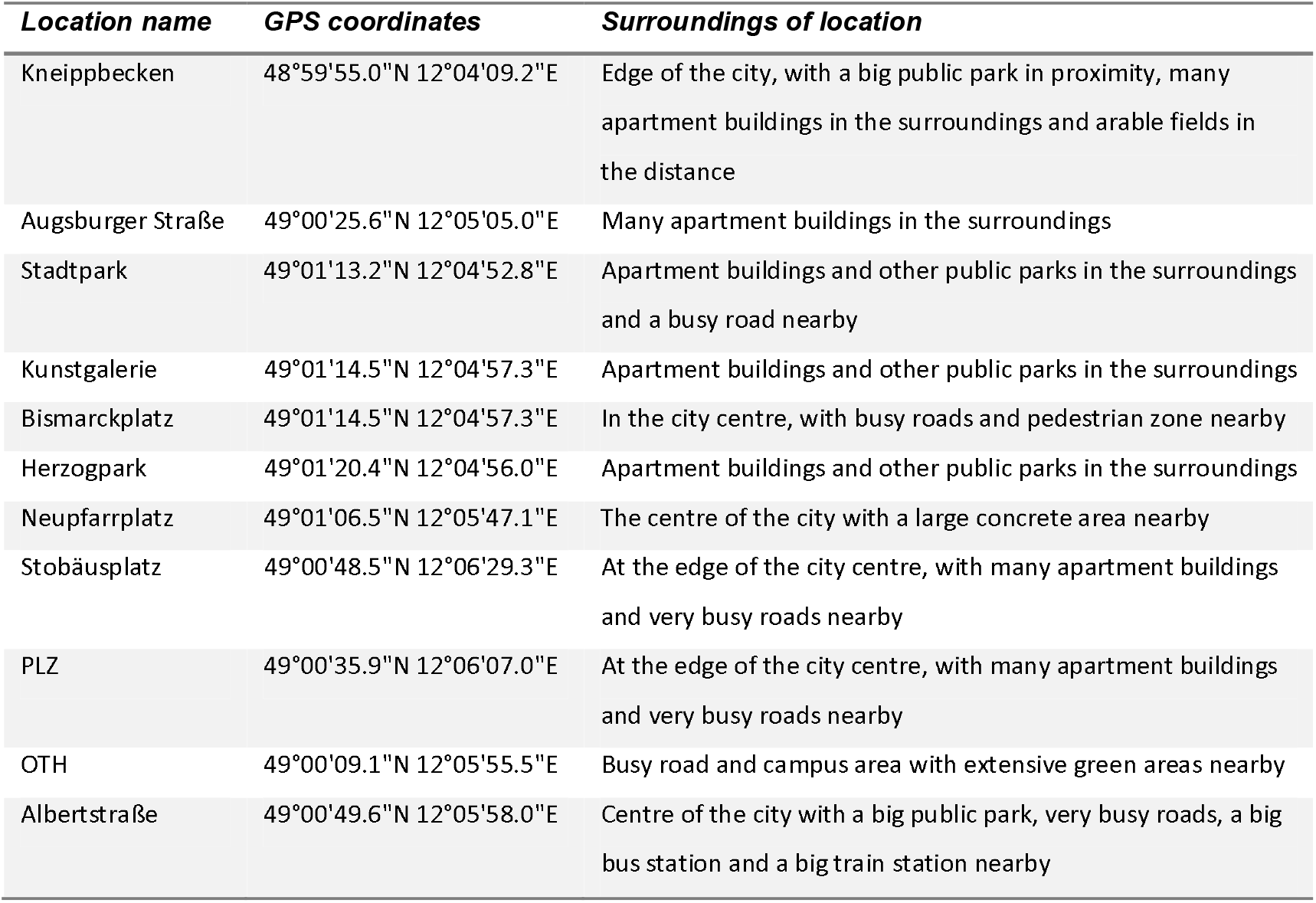
The study locations with GPS coordinates and a brief description.

In each location, a patch of each plant type available was examined. To do this we carried out a standardised observation following Kalaman et al. (2022) in which a 40 x 60cm quadrat was placed on the planted area, and all insects in the area or entering the area over 1 minute were recorded. We hoped that such an extended observation period, rather than the “snapshot” method deployed in other studies (Garbuzov et al. 2015; Palmersheim et al. 2022), would allow us to search more closely for smaller or more inconspicuous insects, such as ants or small bees. We assigned recorded insects to one of the following morphotypes: honey bees, wild bees, bumble bees, hover flies, other flies, butterflies, wasps, true bugs, and ants. After the observation we took a digital photograph of the quadrat, and from this calculated the percentage coverage of the plant type being studied. Based on this, we could calculate the insect observations per m^2^ per minute for each observation.

Observations were carried out in all areas 6 times; 3 in the morning and 3 time in the evening on different days, leading to at least 6 observations minimum per plant type The planting (patch) size of each recorded area was noted, although previous studies indicated that the number of insect visitors per unit area is not affected by patch size, at least within the range of patch sizes typically found within gardens (Garbuzov et al. 2015).

Statistical analysis was carried out in R (R Core Team 2023) using the glmmTMB package (Magnusson et al. 2020) to perform mixed-effect models. Post-hoc pairwise comparisons were carried out using the emmeans package (Lenth et al. 2020). A complete description of the statistical analysis procedure, including all code and statistical output, is provided in supplement S1.

## Results

The complete dataset on which this study is based is provided in supplement S2.

In total, we counted 843 insect visitations over 24 different plant types, over a total of c. 80 hours of data collection. The four most common insect groups were honey bees, wild bees, flies (excluding hoverflies) and ants (figure 1)

**Figure 1).**
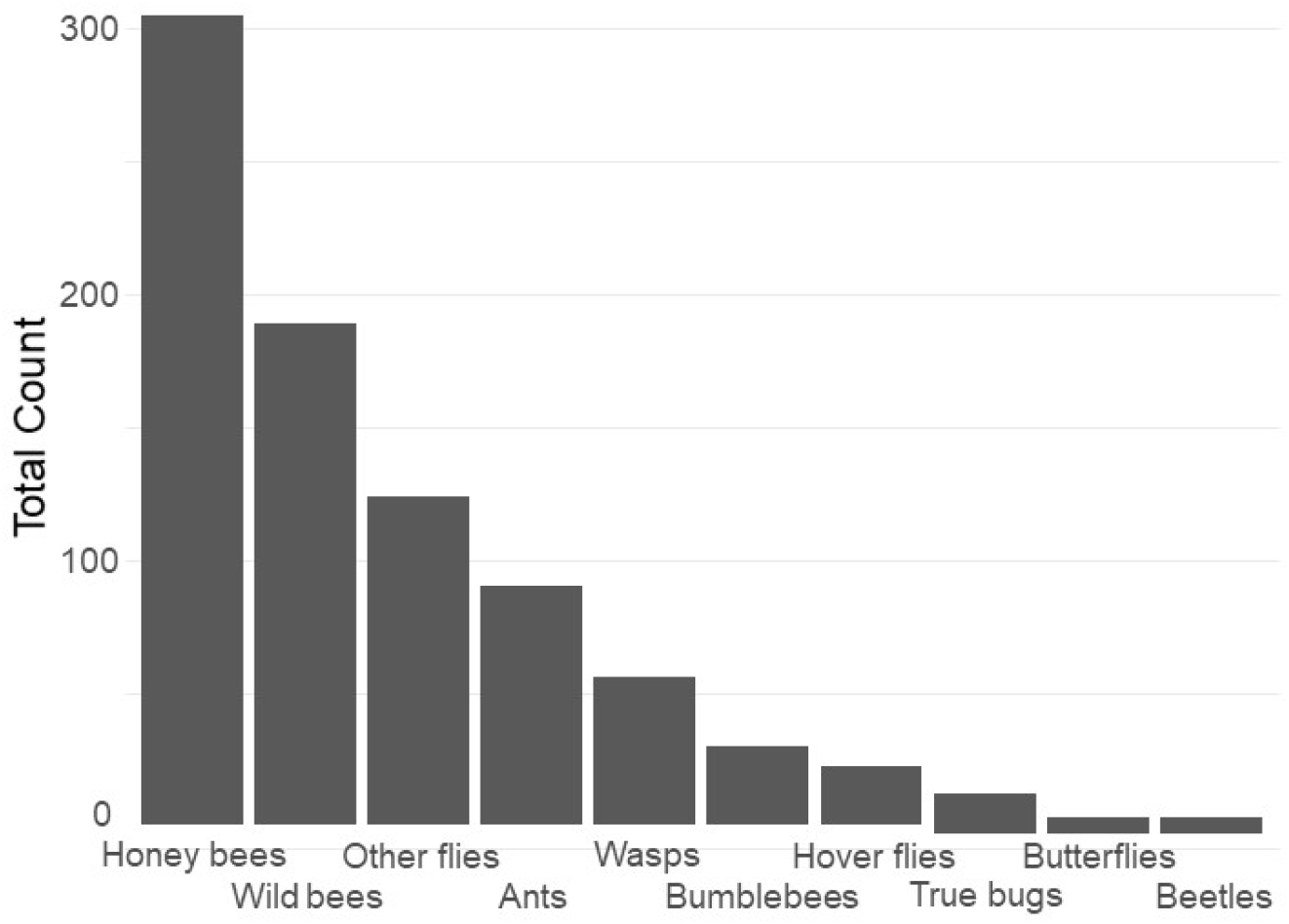
Contribution of each insect group to the total insect count The different plants vary very widely in their visitation rate per m^2^ ground cover (figure 2, right), with Blue Eryngo (*Erynginum planum* “Blaue Zwerg”) having by far the highest visitation rate (mean 5.5 insects min/ m^2^) and French Hydrangea (*Hydrangea macrophylla*) having the lowest (0.041 insects min/ m^2^). However, as Blue Eryngo was so rare in the sample (one location, 6 observations), this measurement is not reliable. The most visited well-sampled plant type was Russian sage (*Salvia yangii*), sampled at 3 locations with 18 observations, which had a visitation rate of 1.61 insects min/ m^2^ . As such, taken at face value, there was a 130-fold difference between the most and least visited plant type, and even considering only the more reliable measurements, there is a 40-fold difference between the most and least visited plant types. Plants differ significantly in their overall attractiveness (glmmTMB, *χ*^2^= 89.06, P < 0.0001). We refrain from individually comparing pairs of plants for brevity – the entire dataset is provided in supplement S1.

As well as being differently attractive, different plant types also attracted a different profile of insects (figure 2 left). For example, while *Salvia yangii* and *Aster ageratoides* show almost identical overall attractiveness in terms of insect visitations per m^2^ min^-1^(figure 2, *Salvia yangii* is dominated by honey bees, while wild bees are the most common visitors to *Aster ageratoides*.

**Figure 2).**
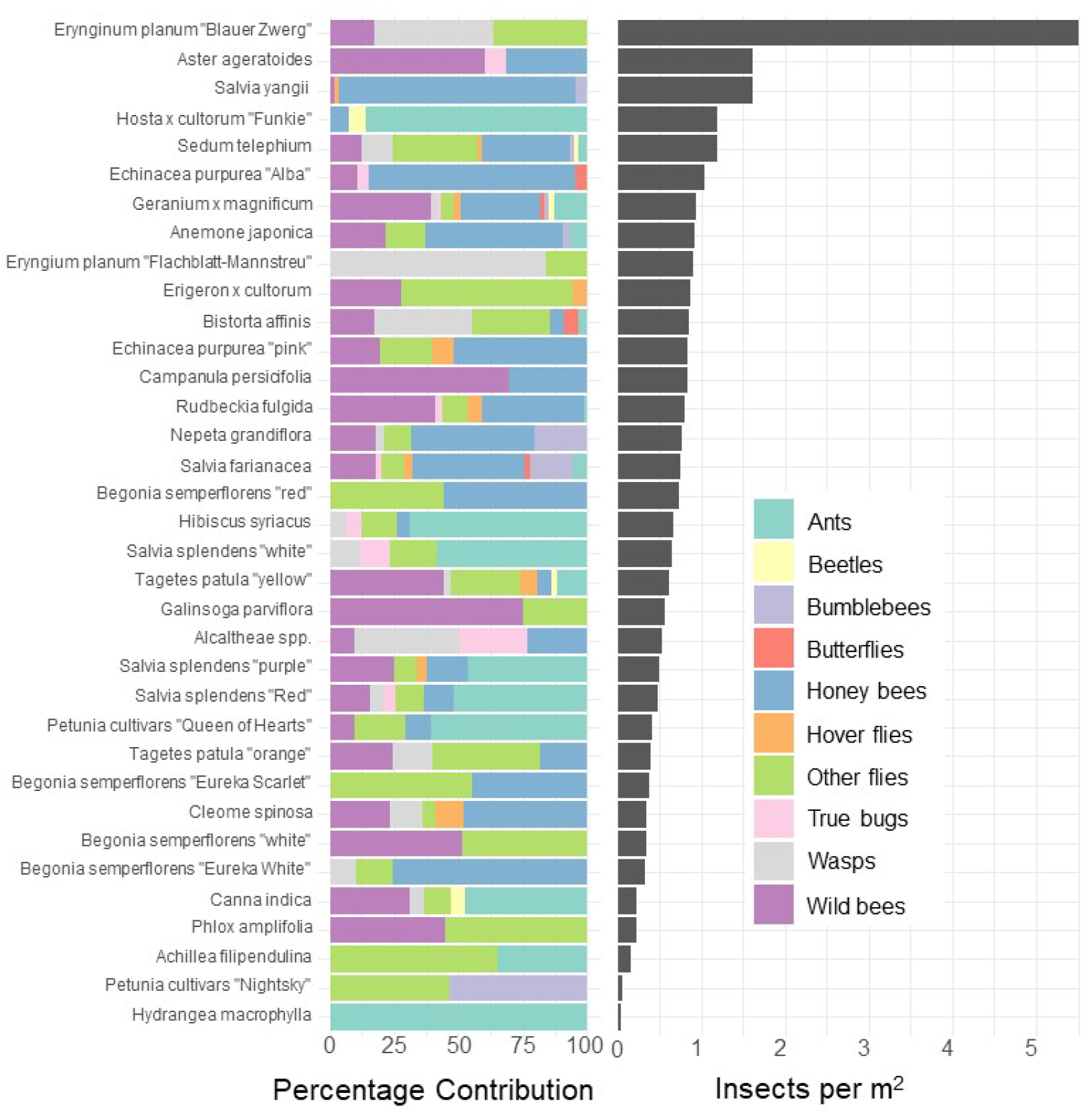
Left: Percentage contribution of each insect type to total visitations for each plant type. Right: Mean visitation rate of all insects to all cultivates, normalised to a rate per m^2^. Note that some plants, such as Blue Eryngo (*Erynginum planum* “Blaue Zwerg”), were not commonly planted, so may not be a reliable estimate.

In order to reduce dimensionality and focus on ecologically relevant topics, we examined the attractiveness of the different cultivars to honey bees and wild bees. Cultivars vary strongly in their attractiveness of honey bees and wild bees (fig 3). The cultivar which attracted the most wild bees per m^2^ was Aster ageratoides followed closely by Erynginum planum “Blaue Zwerg”.

**Figure 3).**
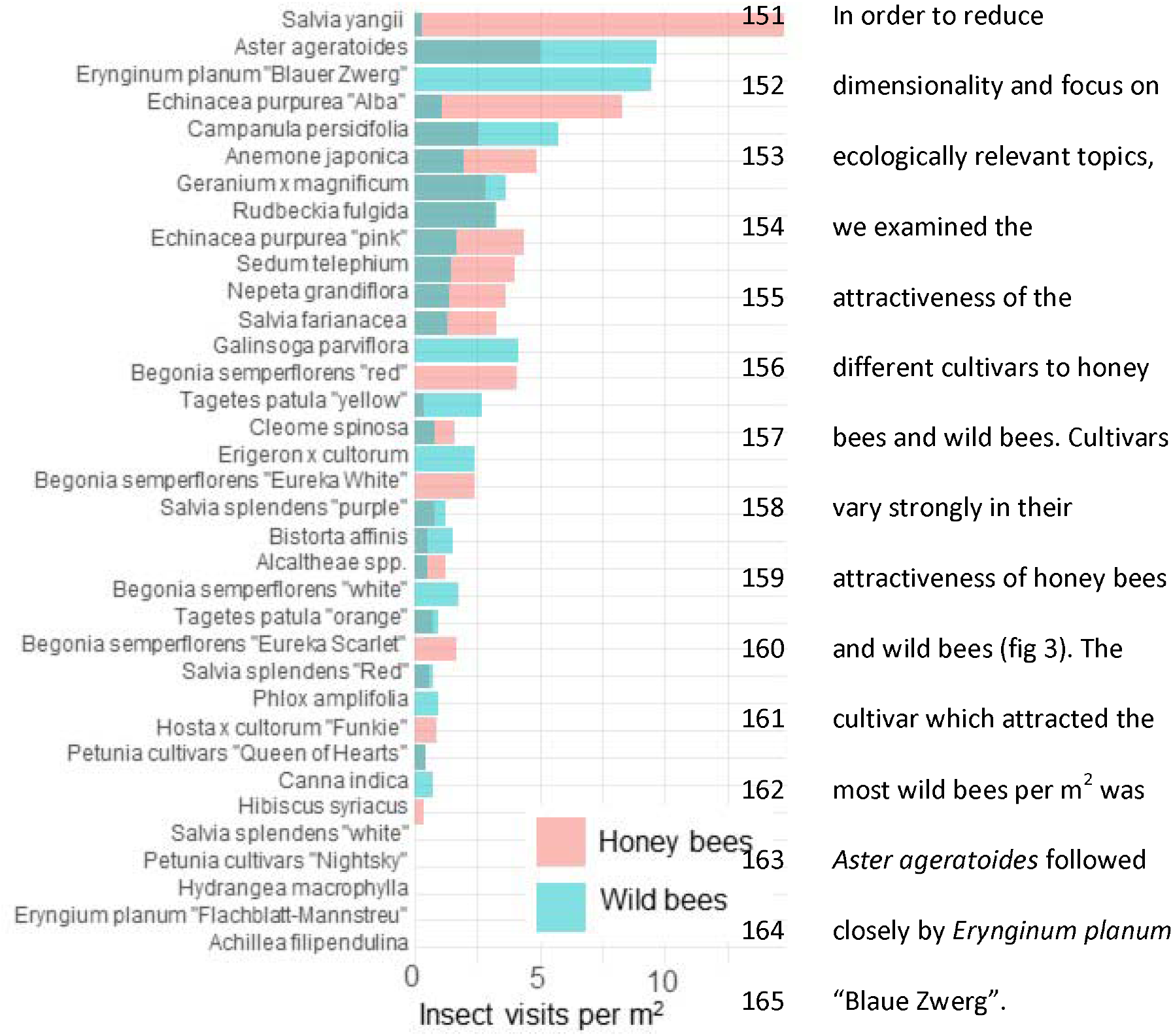
average visitation rate per M^2^ of different plants by honey bees and wild bees.

To further reduce dimensionality, and to allow for model convergence, we only consider the top 6 most attractive flowers in terms of mean insect visits per m^2^ for honey bees and wild bees: Salvia yangii, *Aster ageratoides, Erynginum planum* “Blaue Zwerg”, *Echinacea purpurea* “Alba”, *Campanula persicifolia* and *Anemone japonica*. There was a significant interaction between cultivar and insect group (GLMM, *χ*^2^ = 19.5, df = 5, P = 0.0015). Post-hoc contrast analysis found that more honey bees than wild bees visited *Echinacea purpura “Alba”* (P = 0.083) and *Salvia yangii* (P=0.004) while none of the top 6 most attractive cultivars were visited more often by wild bees. However, it is worth noting that no honey bees, but some wild bees, visited the *Erynginum planum* “Blaue Zwerg”, and twice as many wild bees and honey bees visited *Aster ageratoides*. Unfortunately, these plants were too uncommon in the dataset to draw firm conclusions from. We note that these two cultivars were not found in the same location.

## Discussion

We found enormous (up to 40-130 fold) differences of attractiveness between ornamental plants, mirroring previous studies (Garbuzov et al. 2015; Marquardt et al. 2021a, b; Palmersheim et al. 2022). As all plants surveyed were being planted by the municipality already, this suggests that major improvements to the supply of forage for urban pollinators can be affected rapidly with little effort or additional costs.

Notable in their relative absence from our dataset were bumble bees (3.6% of visitations) and butterflies (0.7%). It is likely that the lack of bumble bees was driven by them not being present: many of the flowers surveyed have previously been reported to be highly attractive to bumblebees, such as *Echinacea purpurea* (Rollings and Goulson 2019). The absence of bumble bees may well be due to the time of year of the survey – many of the local bumble bee species are coming to the end of their annual life-cycle by late summer. A drought in June and July of that year may have also had an effect. However, the urban parks within Regensburg may also not offer a suitable habitat for them, although higher levels of bumble bee visitations were reported in other surveys of urban pollinators (Lowenstein et al. 2019; Honchar and Gnatiuk 2020; Marquardt et al. 2021a). The lack of butterflies may also be explained by the season, although it is likely that the selection of flowers offered were also not especially attractive to honey bees. Plants such as Buddleia or Lantana are especially attractive to butterflies (Shackleton and Ratnieks 2016).

Insect groups and species vary widely in their adaptability to urban environments. Beetles, flies, lepidopterans and solitary bees have been previously reported to be highly sensitive to increasing urbanisation, while generalist insects such as *Bombus terrestris* and *Apis melifera* are not (Bergerot et al. 2011; Mulieri et al. 2011; Geslin et al. 2013; Verboven et al. 2014; Theodorou et al. 2020). This only partially overlaps with our findings (see table 1), and indeed other studies, such as Marquardt et al. (2021a) also report a very large proportion of solitary bees on urban flower, though they also found bumble bees to be rather common. Many wild bees are also generalists, so well adapted to urban environments. Rollings and Goulson (2019) report generally few solitary bees in a plant nursery embedded in a rural environment, except very late in the season (mid-October) when solitary bees suddenly become very common. It is likely that plant types presence, macro-environment, and time of year all strongly affect the number and distribution of attracted insects, highlighting the importance of collecting local data on plant attractiveness.

Our results demonstrate that the current urban plantings do not support a diverse pollinator community. The majority of plants are attractive to honey and wild bees, but not attractive to most other groups, especially butterflies and beetles. At first glance this is disappointed, especially given the presence of locally extremely rare beetles in the city of Regensburg, such as *Nematodes filum* (Regensburg conservation office, unpublished data). However, it is not clear whether broad conservation of all pollinator groups, or rare species, should be the goal of urban planners. Marquardt et al. (2021a) note that “Pollinator conservation strategies in urban areas differ highly in comparison to the strategies in natural or near-natural environments. In cities, the aim is to increase the overall abundance and diversity of pollinator communities, rather than focusing on rare or endangered species.” An achievable aim may be to support wild bees as a class, while possibly avoiding planting plants which are predominantly attractive to honey bees. Honey bees, as a commercially-bred species, are not a conservation priority, and may indeed compete with other pollinators for resources (although direct measurements of impact are often lacking) (Steffan-Dewenter and Tscharntke 2000; Paini 2004; Wojcik et al. 2018), or potentially act as a source of infection (Fürst et al. 2014; Graystock et al. 2016) (but see e.g. (Dolezal et al. 2016; Mallinger et al. 2017)). Bumble bee colonies in close proximity to managed honey bee gives showed longer foraging bout durations (Theodorou et al. 2022). Especially interesting are the decorative thistles, such as the Blue Eryngo, which were very attractive to wild bees, wasp, and flies, but were never visited by honey bees during our observations.

A somewhat surprising finding was the relatively common presence of ants on flowers (10.7% of insects). While surveys of flower attraction to different insect groups are growing in number, ants are very rarely mentioned. It is not clear whether this is because they are ignored, since they rarely act as pollinators (Rostás and Tautz 2011) and many indeed repel pollinators (Junker et al. 2007), or whether they are not noted. Other smaller insect groups, such as thrips, are also hardly reported. It is possible that the underreporting of small, non- or rarely flying insects is due to the predominance of snapshot surveying, which involves noting what can be seen at a glance. We suspect this will tend to bias samples towards big and mobile groups such as honey bees, bumble bees, and butterflies, and away from more cryptic groups such as small wild bees, ants, and thrips. The approach we took, of surveying a patch for a set period of time, may be less prone to such biases, but a systematic comparison of both survey types would be a worthwhile endeavour.

It is important to note that the current study, as almost all other such studies, assumes that plants which are more highly visited are more beneficial to pollinators. This may not always be true. Artificially-bred cultivars could act as cognitive traps, providing a super-stimulus of colour or odour without providing a large reward, or only providing an unbalanced reward. While there is a correlation between nutritional offering and attractiveness (Comba et al. 1999; Corbet et al. 2001; SChapman et al. 2023), little information exists about the nectar or pollen productivity of most ornamental plants, nor of the nutritional content of the nectar and pollen offered.

This study demonstrated that municipally-planted decorative annual flowers vary enormously both in their absolute attractiveness to insects, and in which groups they attract. This suggests that evidence-based changes in municipally-planted flowers could achieve a very rapid improvement in how suitable urban environments are for pollinators, at a very modest financial and organisational expenditure. Indeed, based on the evidence presented in the current study, we have begun a collaboration with the Regensburg municipality to first collect more targeted information about the most promising decorative plants, and ultimately to focus planting on plant species attractive to the local insect groups we wish to promote. However, while promising, some caution is warranted; while this and many studies focus solely on nutritional provision by flowers, not all resources pollinators require come from flowers. Aphids on trees are a very important source of carbohydrates for many insects, and many insects require non-food resources such as resin or nesting sites, which are better provided by trees and bushes (Requier and Leonhardt 2020). Forage for the larval stages of pollinators is also an important aspect to keep in mind. Improving the attractiveness of municipally-planted flowers is a relatively quick and easy approach, with potentially large rewards, but it can only be one part of a larger programme for supporting urban biodiversity.

## Acknowledgements

Many thanks to Daniela Feuerer for help with cultivar identification. TJC was funded by the Deutsche Forschungsgemeinschaft (DFG, German Research Foundation) – project number 462101190.

