## supplement S2 for "High variability in the attractiveness of municipally-planted decorative plants to pollinators"

2024-01-19

#### packages

```
library (ggplot2)
library(readxl)
library(forcats)
library (glmmTMB)
library (dplyr)
library (DHARMa)
library (multcomp)
library(emmeans)
library (eoffice)
library(gridExtra)
library(car)
```

### input data, subsetting

```
data <- read_excel("municipal_pollinator_data_merged_bumblebees.xlsx",
    sheet = "global_database_goodnames")

ants <- subset (data, insect_group == "Ants")
flies <- subset (data, insect_group == "Other flies")
honeybees <- subset (data, insect_group == "Honey bees")
wasps <- subset (data, insect_group == "Wasps")
wildbees <- subset (data, insect_group == "Wild bees")

bees_and_wildbees <- rbind(honeybees, wildbees)
bees_wildbees_flies_wasps <- rbind(honeybees, wildbees, flies, wasps)
```

### plots

#### plot of all insect visitation rates for all cultivars

First some summary statistics: mean visitation rate for each cultivar

```
mean_insectsM2 <- data %>%
  group_by(cultivar) %>%
  summarise(mean_insects_m2 = mean(insects_m2_full_coverage, na.rm = TRUE))

mean_insectsM2
```

```
## # A tibble: 35 x 2
##    cultivar                                   mean_insects_m2
##    <chr>                                                <dbl>
##  1 "Achillea filipendulina"                             0.160
##  2 "Alcaltheae spp."                                    0.529
##  3 "Anemone japonica"                                   0.914
##  4 "Aster ageratoides"                                  1.61
##  5 "Begonia semperflorens \"Eureka Scarlet\""           0.377
##  6 "Begonia semperflorens \"Eureka White\""             0.318
##  7 "Begonia semperflorens \"red\""                      0.735
##  8 "Begonia semperflorens \"white\""                    0.334
##  9 "Bistorta affinis"                                   0.857
## 10 "Campanula persicifolia"                             0.828
## # ... with 25 more rows
```

now some descriptive plots

```
# mean bar plot
all_insects_by_cultivar_plot <- ggplot (data, aes(x = reorder(cultivar,insects_m2_full_coverage), y = i
 scale_y_continuous(expand = c(0, 0)) + # forces X axis to 0
  coord_cartesian(ylim = c(0, 6)) +
  stat_summary(fun.y = "mean", geom = "bar",  size = 3) +
  theme_minimal() +
  theme(axis.text.x = element_text(angle = 90, hjust = 1,vjust = 0.3)) +
```

```
  labs( x = "cultivar", y = "Insects per m2", title ="all insects")

plot (all_insects_by_cultivar_plot)
```

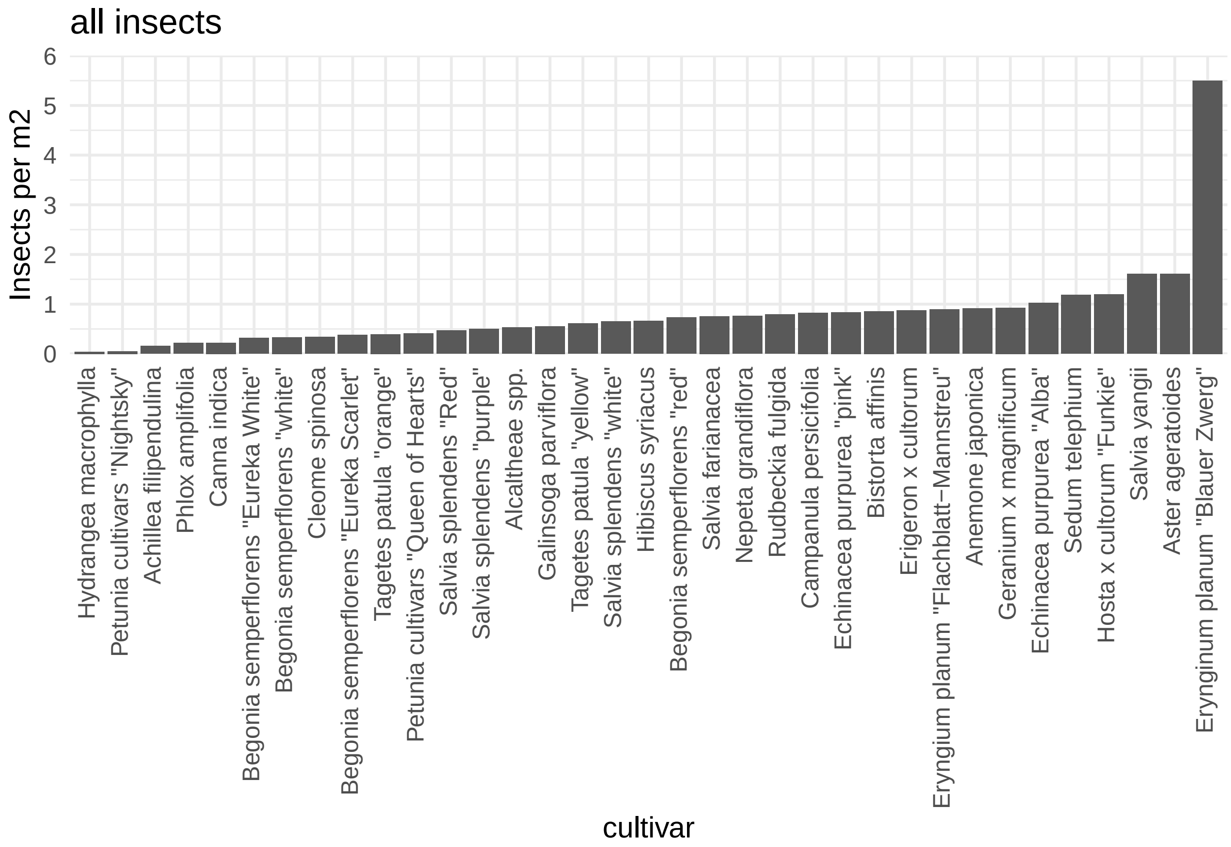

```
#topptx (all_insects_by_cultivar_plot, "fig2_all_insects_by_cultivar_plot.pptx", width = 6, height = 8)
```

Now individual plots of cultivar attractiveness forthe top 4 most common insect groups (honeybees wildbees flies ants wasps)

```
insectplot_facet_common <- ggplot (bees_wildbees_flies_wasps, aes(x = reorder(cultivar,insects_m2_full_
  facet_wrap(~ insect_group, nrow = 4, ncol = 1)+
  scale_y_continuous(expand = c(0, 0)) + # forces X axis to 0, but in this case is overriden by ribbon
  coord_cartesian(ylim = c(0, 26)) +
  stat_summary(fun.y = "mean", geom = "bar",  size = 3, fill = "red") +
  ylab("Average insects per M2") +
  xlab("Cultivar") +
  theme_bw(15) +
  theme(axis.text.x = element_text(angle = 90, hjust = 1,vjust = 0.3))

plot (insectplot_facet_common)
```

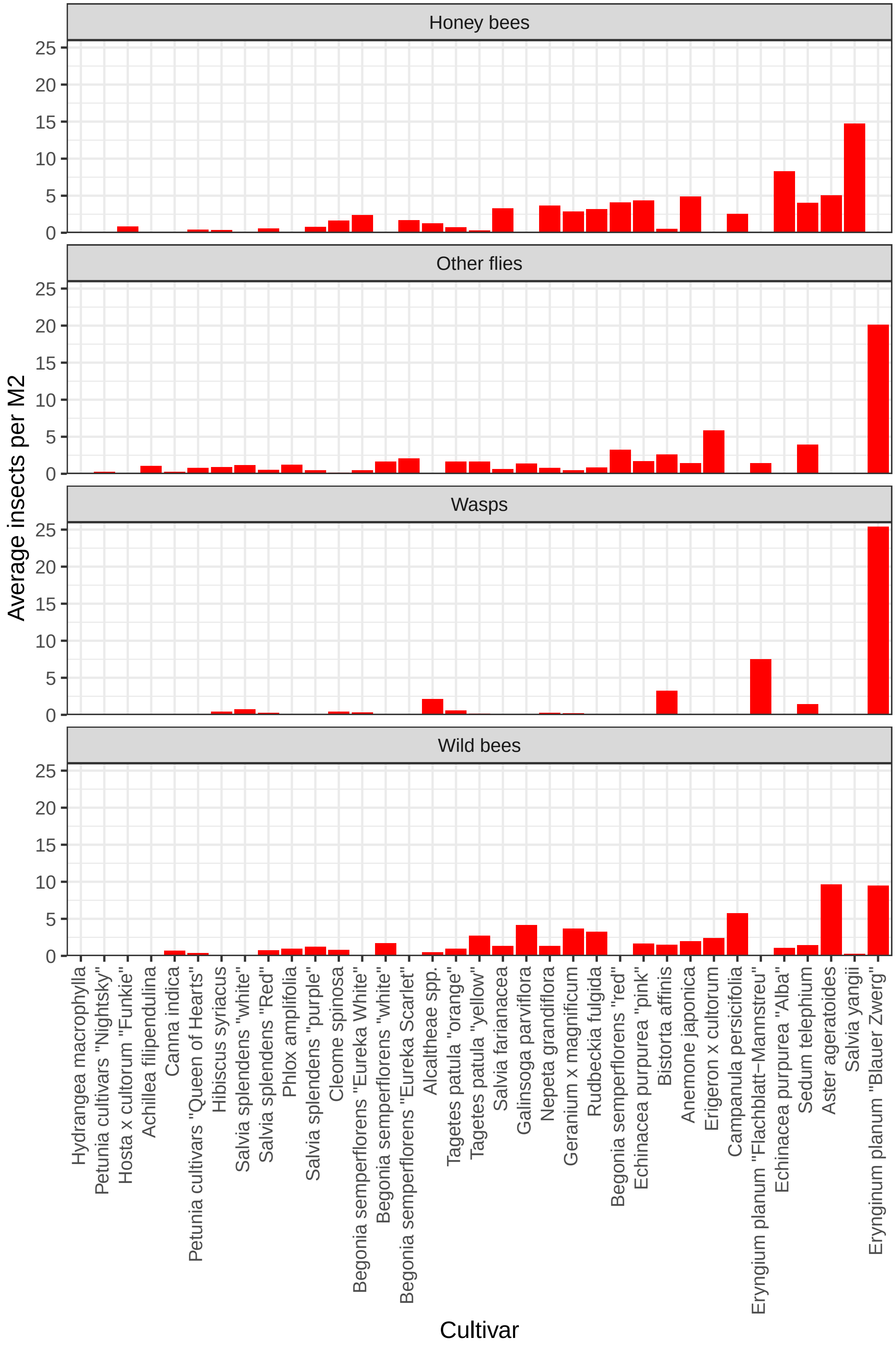

Honey bees
Other flies
Wasps
Wild bees
Average insects per M2
Cultivar
0
5
10
15
20
25
Hydrangea macrophylla
Petunia cultivars "Nightsky"
Hosta x cultorum "Funkie"
Achillea filipendulina
Canna indica
Petunia cultivars "Queen of Hearts"
Hibiscus syriacus
Salvia splendens "white"
Salvia splendens "Red"
Phlox amplifolia
Salvia splendens "purple"
Cleome spinosa
Begonia semperflorens "Eureka White"
Begonia semperflorens "white"
Begonia semperflorens "Eureka Scarlet"
Alcaltheae spp.
Tagetes patula "orange"
Tagetes patula "yellow"
Salvia farianacea
Galinsoga parviflora
Nepeta grandiflora
Geranium x magnificum
Rudbeckia fulgida
Begonia semperflorens "red"
Echinacea purpurea "pink"
Bistorta affinis
Anemone japonica
Erigeron x cultorum
Campanula persicifolia
Eryngium planum "Flachblatt–Mannstreu"
Echinacea purpurea "Alba"
Sedum telephium
Aster ageratoides
Salvia yangii
Eryngium planum "Blauer Zwerg"

#### Total contributions of cultivars and insect groups, in terms of insects per m2

##### Contribution of each insect group to total

First some summary statistics

```
data_summary <- data %>%
  group_by(insect_group) %>%
  summarise(total_count = sum(count)) %>%
  mutate(percentage = (total_count / sum(total_count)) * 100) %>%
  arrange(desc(percentage))

data_summary
```

```
## # A tibble: 10 x 3
##    insect_group total_count percentage
##    <chr>              <dbl>      <dbl>
##  1 Honey bees           305     36.2
##  2 Wild bees            189     22.4
##  3 Other flies          124     14.7
##  4 Ants                  90     10.7
##  5 Wasps                 56      6.64
##  6 Bumblebees            30      3.56
##  7 Hover flies           22      2.61
##  8 True bugs             15      1.78
##  9 Beetles                6      0.712
## 10 Butterflies            6      0.712
```

first just one bar chart showing contribution of all insect groups

```
# Calculate the total count for each insect group
total_counts <- data %>%
  group_by(insect_group) %>%
  summarise(total_count = sum(count)) %>%
  arrange(desc(total_count))  # Order by total count in descending order

# Create a bar chart
insect_contribution_plot <- ggplot(total_counts, aes(x = reorder(insect_group, -total_count), y = total_
  geom_bar(stat = "identity") +
  labs( x = "Insect Group",
       y = "Total Count") +
  theme_minimal()

plot (insect_contribution_plot)
```

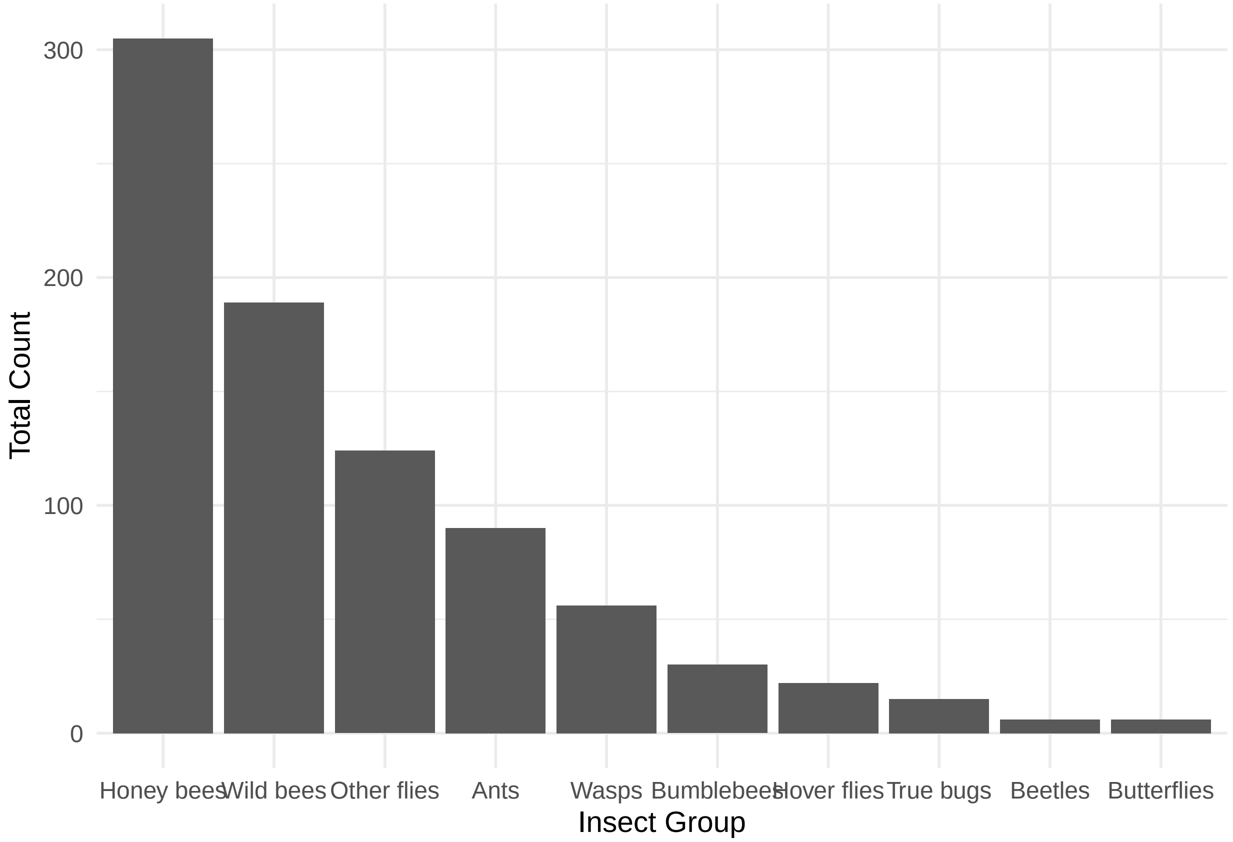

```
#topptx (insect_contribution_plot, "insect_contribution_plot.pptx", width = 6, height = 6)
```

###### Insect group visitation proportion by cultivar

This helps us see who the visitors are to each cultivar

```
# Calculate the percentage contribution of 'insects_m2_full_coverage' for each 'insect_group' within ea
data_percent <- data %>%
  group_by(cultivar, insect_group) %>%
  summarise(total_insects_m2 = sum(insects_m2_full_coverage),
            original_values = mean(insects_m2_full_coverage)) %>%
  group_by(cultivar) %>%
  mutate(percentage = total_insects_m2 / sum(total_insects_m2) * 100)
```

```
## 'summarise()' has grouped output by 'cultivar'. You can override using the '.groups' argument.
```

```
# Create a stacked bar chart showing the percentage contribution
filledstackedplot_cultivar <- ggplot(data_percent, aes(x = reorder(cultivar,original_values), y = percen
  geom_bar(stat = "identity") +
  labs(title = "Percentage Contribution of Insect Groups to Cultivars",
       x = "Cultivar", y = "Percentage Contribution (%)") +
  scale_fill_brewer(palette = "Set3") +  # Choose a color palette
  theme_minimal() +
  theme(axis.text.x = element_text(angle = 90, hjust = 1,vjust = 0.3))
```

```
plot (filledstackedplot_cultivar)
```

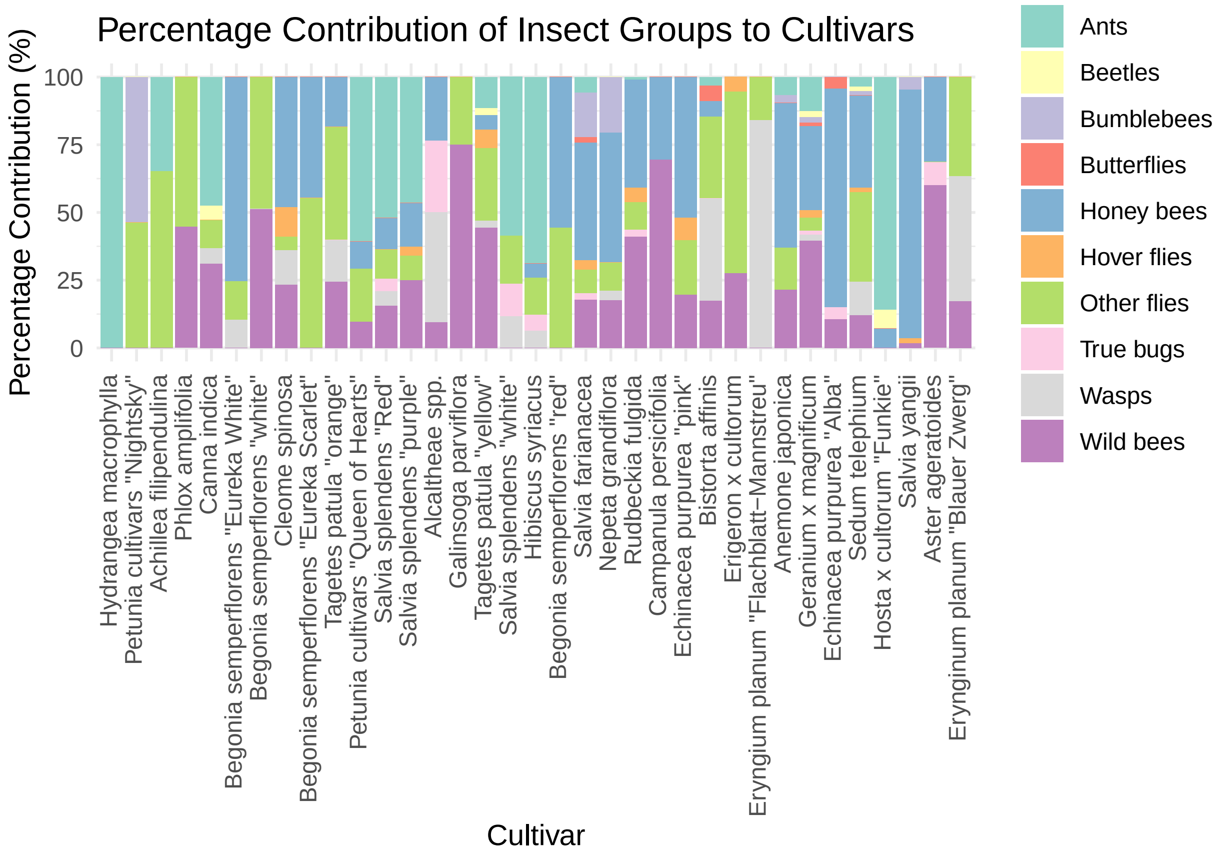

```
#topptx (filledstackedplot_cultivar, "fig3_filledstackedplot_cultivar.pptx", width = 8, height = 6)
```

##### total planted area size vs insect visitation rate

Various publications claim that there is no relationship between planted area size and insect visitation rates per unit area. I will quickly confirm visually.

```
ggplot (data, aes(x = total_area_cultivars, y = insects_m2_full_coverage)) +
geom_jitter(width = 10, height = 1, alpha = 0.3) +
  geom_smooth(method = "lm") +
  theme_bw(15) +
  theme(axis.text.x = element_text(angle = 90, hjust = 1)) +
  labs( x = "total area of planted plants in location", y = "Insects per m2")
```

```
## `geom_smooth()` using formula 'y ~ x'
```

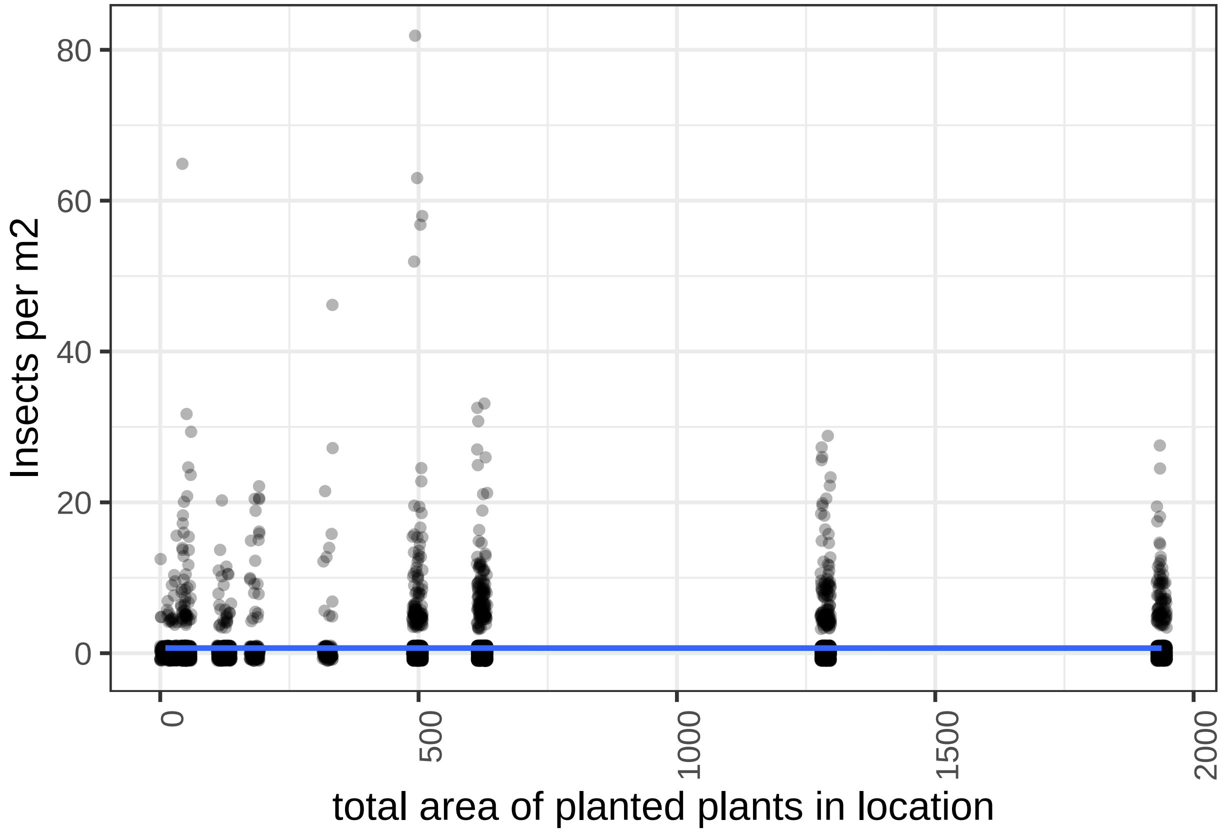

Confirmed. No apparent effect. Naturally, this is also heavily skewed by the zero inflation, but certainly nothing obvious here. A quick statistical analysis to be found in the stats part.

##### Wild bees and honeybees - plots

To reduce dimensionality and persue clear questions, here we are focussing in on just wild bees and honeybees. Is there an overlap between wild bees and honeybees? What plants do wild bees like but honeybees don't?

```
fig4_honey_wildbees <- ggplot (bees_and_wildbees, aes(x = reorder(cultivar,insects_m2_full_coverage), y
  scale_y_continuous(expand = c(0, 0)) + # forces X axis to 0, but in this case is overriden by ribbon
  coord_cartesian(ylim = c(0, 20)) +
  stat_summary(fun.y = "mean", geom = "bar", shape = 23, size = 3, alpha = 0.5) +
  theme_minimal() +
  theme(axis.text.x = element_text(angle = 90, hjust = 1, vjust = 0.3)) +  # hjust and vjust nudge the 
  labs( x = "cultivar", y = "Insects per m2", title ="mean M2 honey and wildbees")

plot (fig4_honey_wildbees)
```

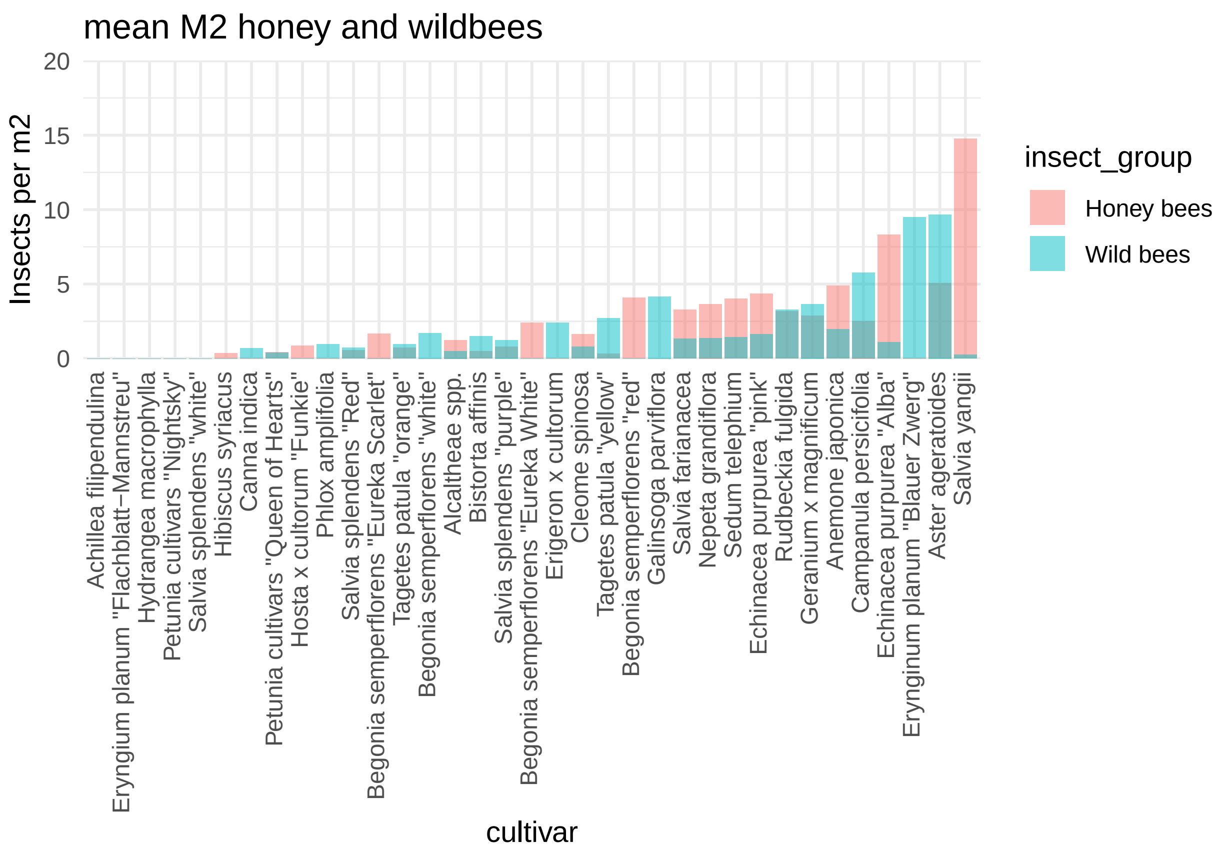

```
#topptx (fig4_honey_wildbees, "fig4_wildvshoney.pptx", width = 6, height = 6)
```

some descriptive states here:

sum of count

```
# Calculate the sum of "count" for each combination of "cultivar" and "insect_group"
bees_and_wildbees %>%
  group_by(cultivar, insect_group) %>%
  summarise(total_count = sum(count))
```

```
## `summarise()` has grouped output by 'cultivar'. You can override using the `.groups` argument.
```

```
## # A tibble: 70 x 3
## # Groups:   cultivar [35]
##    cultivar                                    insect_group total_count
##    <chr>                                       <chr>              <dbl>
##  1 "Achillea filipendulina"                    Honey bees             0
##  2 "Achillea filipendulina"                    Wild bees              0
##  3 "Alcaltheae spp."                           Honey bees             2
##  4 "Alcaltheae spp."                           Wild bees              1
##  5 "Anemone japonica"                          Honey bees            27
##  6 "Anemone japonica"                          Wild bees             12
##  7 "Aster ageratoides"                         Honey bees             7
##  8 "Aster ageratoides"                         Wild bees             13
```

```
##  9 "Begonia semperflorens \"Eureka Scarlet\"" Honey bees             4
## 10 "Begonia semperflorens \"Eureka Scarlet\"" Wild bees              0
## # ... with 60 more rows
```

```
bees_and_wildbees %>%
  group_by(insect_group) %>%
  summarise(total_count = sum(count))
```

```
## # A tibble: 2 x 2
##   insect_group total_count
##   <chr>              <dbl>
## 1 Honey bees           305
## 2 Wild bees            189
```

#### statistical analysis

Questions we want to ask: - do cultivars differ in attractiveness overall? - do cultivears differ in attractiveness for specific target groups? - does total patch area affect insect visitation rates - then do pairwise comparisons - which cultivars has more wild bees than honeybees?

##### do cultivars differ in attractiveness?

```
m1 <- glmmTMB(insects_m2_full_coverage ~ cultivar
              + (1 | location),
              ziformula=~1,
              family=nbinom1,
              data = data)
```

looking at results, testing model fit

```
#simulateResiduals(m1, n = 500, plot = T) #dharma

adjX <- glht(m1, alternative = "two.sided") # multcomp

summary(adjX, test = adjusted("BH"))
```

```
##
##   Simultaneous Tests for General Linear Hypotheses
##
## Fit: glmmTMB(formula = insects_m2_full_coverage ~ cultivar + (1 |
##     location), data = data, family = nbinom1, ziformula = ~1,
##     dispformula = ~1)
##
## Linear Hypotheses:
##                                                      Estimate Std. Error z value
## (Intercept) == 0                                      1.69469    0.60692   2.792
## cultivarAlcaltheae spp. == 0                          0.14181    0.64412   0.220
## cultivarAnemone japonica == 0                         0.59901    0.61460   0.975
```

```
## cultivarAster ageratoides == 0                      0.40294    0.62316   0.647
## cultivarBegonia semperflorens "Eureka Scarlet" == 0  0.53668    0.65169   0.824
## cultivarBegonia semperflorens "Eureka White" == 0   -0.04650    0.62876  -0.074
## cultivarBegonia semperflorens "red" == 0             0.23150    0.72878   0.318
## cultivarBegonia semperflorens "white" == 0           0.25822    0.68265   0.378
## cultivarBistorta affinis == 0                        0.37179    0.62214   0.598
## cultivarCampanula persicifolia == 0                  0.84611    0.66669   1.269
## cultivarCanna indica == 0                            0.16360    0.64022   0.256
## cultivarCleome spinosa == 0                         -0.01248    0.62178  -0.020
## cultivarEchinacea purpurea "Alba" == 0               0.45831    0.62413   0.734
## cultivarEchinacea purpurea "pink" == 0               0.50941    0.61761   0.825
## cultivarErigeron x cultorum == 0                     0.75820    0.63069   1.202
## cultivarErynginum planum "Blauer Zwerg" == 0         2.40088    0.61694   3.892
## cultivarEryngium planum "Flachblatt-Mannstreu" == 0  1.59038    0.64522   2.465
## cultivarGalinsoga parviflora == 0                    0.26557    0.73107   0.363
## cultivarGeranium x magnificum == 0                   0.40775    0.60995   0.669
## cultivarHibiscus syriacus == 0                       0.32816    0.63490   0.517
## cultivarHosta x cultorum "Funkie" == 0               0.82405    0.64045   1.287
## cultivarHydrangea macrophylla == 0                  -3.18657    1.22588  -2.599
## cultivarNepeta grandiflora == 0                      0.22393    0.62556   0.358
## cultivarPetunia cultivars "Nightsky" == 0           -3.23855    0.97183  -3.332
## cultivarPetunia cultivars "Queen of Hearts" == 0     0.46119    0.66103   0.698
## cultivarPhlox amplifolia == 0                        0.56841    0.69933   0.813
## cultivarRudbeckia fulgida == 0                       0.39484    0.60951   0.648
## cultivarSalvia farianacea == 0                       0.31851    0.61010   0.522
## cultivarSalvia splendens "purple" == 0               0.70736    0.62108   1.139
## cultivarSalvia splendens "Red" == 0                  0.45030    0.61741   0.729
## cultivarSalvia splendens "white" == 0                0.33364    0.65074   0.513
## cultivarSalvia yangii == 0                           1.00351    0.61436   1.633
## cultivarSedum telephium == 0                         0.43070    0.61380   0.702
## cultivarTagetes patula "orange" == 0                 0.53862    0.63136   0.853
## cultivarTagetes patula "yellow" == 0                 0.46713    0.62105   0.752
##                                                     Pr(>|z|)
## (Intercept) == 0                                     0.06106 .
## cultivarAlcaltheae spp. == 0                         0.87579
## cultivarAnemone japonica == 0                        0.78809
## cultivarAster ageratoides == 0                       0.78809
## cultivarBegonia semperflorens "Eureka Scarlet" == 0  0.78809
## cultivarBegonia semperflorens "Eureka White" == 0    0.96872
## cultivarBegonia semperflorens "red" == 0             0.84762
## cultivarBegonia semperflorens "white" == 0           0.84044
## cultivarBistorta affinis == 0                        0.78835
## cultivarCampanula persicifolia == 0                  0.78809
## cultivarCanna indica == 0                            0.87314
## cultivarCleome spinosa == 0                          0.98398
## cultivarEchinacea purpurea "Alba" == 0               0.78809
## cultivarEchinacea purpurea "pink" == 0               0.78809
## cultivarErigeron x cultorum == 0                     0.78809
## cultivarErynginum planum "Blauer Zwerg" == 0         0.00349 **
## cultivarEryngium planum "Flachblatt-Mannstreu" == 0  0.09595 .
## cultivarGalinsoga parviflora == 0                    0.84044
## cultivarGeranium x magnificum == 0                   0.78809
## cultivarHibiscus syriacus == 0                       0.78835
## cultivarHosta x cultorum "Funkie" == 0               0.78809
```

```
## cultivarHydrangea macrophylla == 0                    0.08171 .
## cultivarNepeta grandiflora == 0                       0.84044
## cultivarPetunia cultivars "Nightsky" == 0             0.01507 *
## cultivarPetunia cultivars "Queen of Hearts" == 0      0.78809
## cultivarPhlox amplifolia == 0                         0.78809
## cultivarRudbeckia fulgida == 0                        0.78809
## cultivarSalvia farianacea == 0                        0.78835
## cultivarSalvia splendens "purple" == 0                0.78809
## cultivarSalvia splendens "Red" == 0                   0.78809
## cultivarSalvia splendens "white" == 0                 0.78835
## cultivarSalvia yangii == 0                            0.59723
## cultivarSedum telephium == 0                          0.78809
## cultivarTagetes patula "orange" == 0                  0.78809
## cultivarTagetes patula "yellow" == 0                  0.78809
## ---
## Signif. codes:  0 '***' 0.001 '**' 0.01 '*' 0.05 '.' 0.1 ' ' 1
## (Adjusted p values reported -- BH method)
```

```
#emmeans

meanie <- emmeans(m1, pairwise ~   cultivar)
# print (meanie) # no point plotting all pairwise comparisons for all pairs - we'd have a heck of a lot
plot (meanie)
```

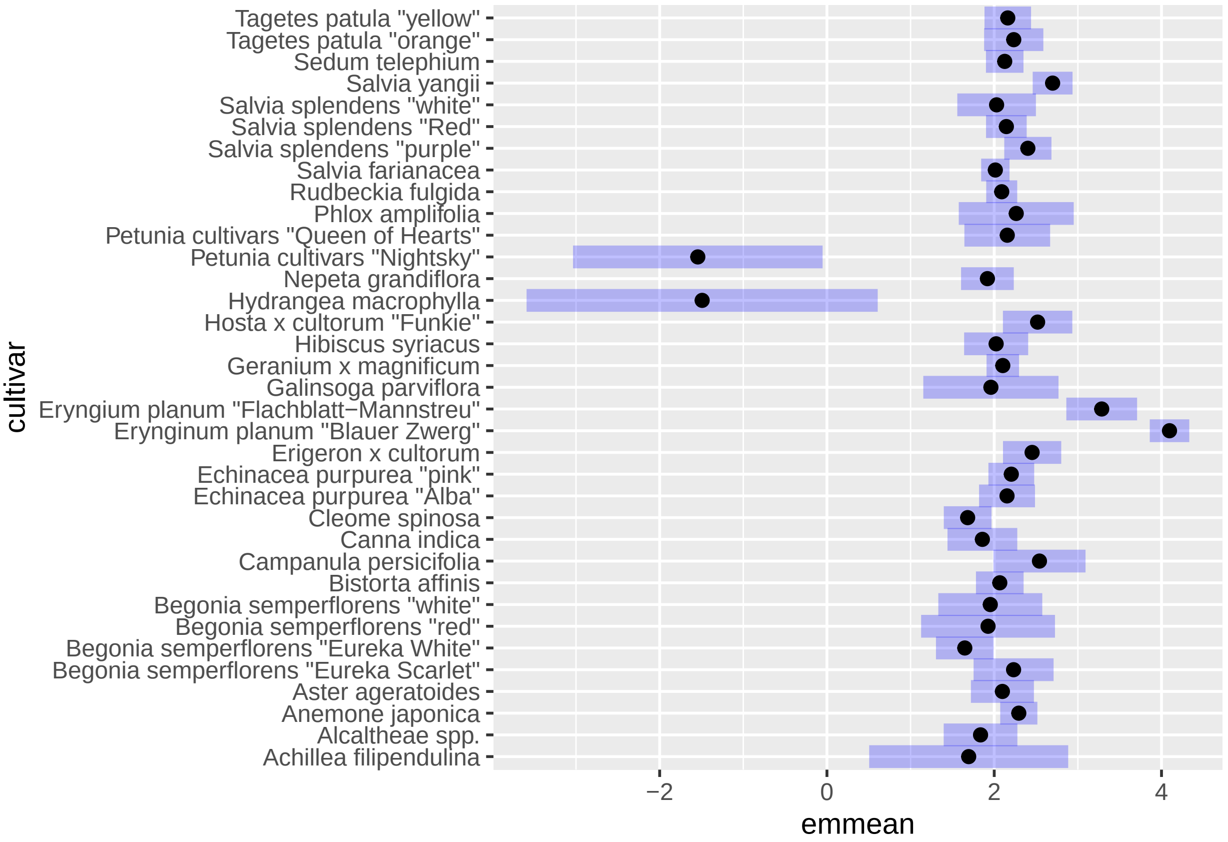

#### Wild bess

Are cultivars differently attractive to wild bees?

```
m2_wildbees <- glmmTMB( # run a GLMMtmb, and put the results in container, in this case called m2_area
  insects_m2_full_coverage ~ cultivar  # try to predict the thing before the ~ by the thing(s) after th
              + (1 | location), # add a random effect (something which might effect the data but we don
              ziformula=~1, # magic to control for zero inflation
              family=nbinom1, # what is the distribution family of your data? gaussian? binomial? Poiss
              data = wildbees) # what data should the model consider?

Anova (m2_wildbees) # this looks at the resulting model. needs the package 'car'
```

```
## Analysis of Deviance Table (Type II Wald chisquare tests)
##
## Response: insects_m2_full_coverage
##           Chisq Df Pr(>Chisq)
## cultivar       34
```

looking at results, testing model fit

```
simulateResiduals(m2_wildbees, n = 500, plot = T) #dharma
```

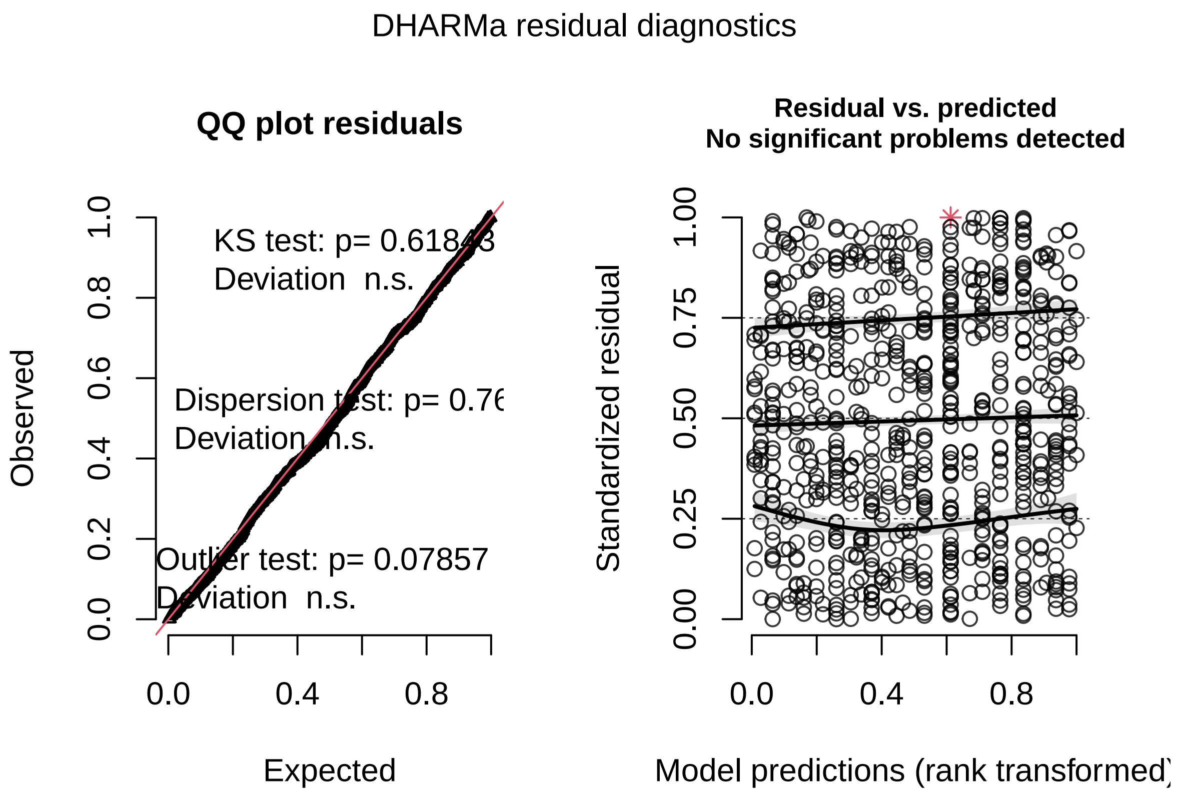

```
## Object of Class DHARMa with simulated residuals based on 500 simulations with refit = FALSE . See ?DI
##
## Scaled residual values: 0.97 0.876 0.1967105 0.916 0.974 0.2334333 0.7300375 0.09202511 0.904 0.956 (
```

```r
adjX <- glht(m2_wildbees, alternative = "two.sided") # multcomp

summary(adjX, test = adjusted("BH"))
```

```
##
##   Simultaneous Tests for General Linear Hypotheses
##
## Fit: glmmTMB(formula = insects_m2_full_coverage ~ cultivar + (1 |
##     location), data = wildbees, family = nbinom1, ziformula = ~1,
##     dispformula = ~1)
##
## Linear Hypotheses:
##                                                        Estimate Std. Error z value
## (Intercept) == 0                                         -15.15         NA      NA
## cultivarAlcaltheae spp. == 0                              16.99         NA      NA
## cultivarAnemone japonica == 0                             17.48         NA      NA
## cultivarAster ageratoides == 0                            17.46         NA      NA
## cultivarBegonia semperflorens "Eureka Scarlet" == 0      -35.81         NA      NA
## cultivarBegonia semperflorens "Eureka White" == 0        -35.19         NA      NA
## cultivarBegonia semperflorens "red" == 0                 -21.51         NA      NA
## cultivarBegonia semperflorens "white" == 0                17.18         NA      NA
## cultivarBistorta affinis == 0                             17.09         NA      NA
## cultivarCampanula persicifolia == 0                       17.67         NA      NA
## cultivarCanna indica == 0                                 16.99         NA      NA
## cultivarCleome spinosa == 0                               16.92         NA      NA
## cultivarEchinacea purpurea "Alba" == 0                    16.96         NA      NA
## cultivarEchinacea purpurea "pink" == 0                    17.12         NA      NA
## cultivarErigeron x cultorum == 0                          17.43         NA      NA
## cultivarEryinginum planum "Blauer Zwerg" == 0             19.21         NA      NA
## cultivarEryngium planum "Flachblatt-Mannstreu" == 0      -34.34         NA      NA
## cultivarGalinsoga parviflora == 0                         17.75         NA      NA
## cultivarGeranium x magnificum == 0                        17.48         NA      NA
## cultivarHibiscus syriacus == 0                           -36.41         NA      NA
## cultivarHosta x cultorum "Funkie" == 0                   -31.88         NA      NA
## cultivarHydrangea macrophylla == 0                       -35.09         NA      NA
## cultivarNepeta grandiflora == 0                           16.94         NA      NA
## cultivarPetunia cultivars "Nightsky" == 0                -33.65         NA      NA
## cultivarPetunia cultivars "Queen of Hearts" == 0          16.59         NA      NA
## cultivarPhlox amplifolia == 0                             16.99         NA      NA
## cultivarRudbeckia fulgida == 0                            17.57         NA      NA
## cultivarSalvia farianacea == 0                            17.33         NA      NA
## cultivarSalvia splendens "purple" == 0                    17.80         NA      NA
## cultivarSalvia splendens "Red" == 0                       17.30         NA      NA
## cultivarSalvia splendens "white" == 0                    -33.22         NA      NA
## cultivarSalvia yangii == 0                                15.37         NA      NA
## cultivarSedum telephium == 0                              17.28         NA      NA
## cultivarTagetes patula "orange" == 0                      17.59         NA      NA
## cultivarTagetes patula "yellow" == 0                      18.05         NA      NA
##                                                        Pr(>|z|)
## (Intercept) == 0                                             NA
## cultivarAlcaltheae spp. == 0                                 NA
## cultivarAnemone japonica == 0                                NA
## cultivarAster ageratoides == 0                               NA
```

```
## cultivarBegonia semperflorens "Eureka Scarlet" == 0       NA
## cultivarBegonia semperflorens "Eureka White" == 0         NA
## cultivarBegonia semperflorens "red" == 0                  NA
## cultivarBegonia semperflorens "white" == 0                NA
## cultivarBistorta affinis == 0                             NA
## cultivarCampanula persicifolia == 0                       NA
## cultivarCanna indica == 0                                 NA
## cultivarCleome spinosa == 0                               NA
## cultivarEchinacea purpurea "Alba" == 0                    NA
## cultivarEchinacea purpurea "pink" == 0                    NA
## cultivarErigeron x cultorum == 0                          NA
## cultivarEryngium planum "Blauer Zwerg" == 0               NA
## cultivarEryngium planum "Flachblatt-Mannstreu" == 0       NA
## cultivarGalinsoga parviflora == 0                         NA
## cultivarGeranium x magnificum == 0                        NA
## cultivarHibiscus syriacus == 0                            NA
## cultivarHosta x cultorum "Funkie" == 0                    NA
## cultivarHydrangea macrophylla == 0                        NA
## cultivarNepeta grandiflora == 0                           NA
## cultivarPetunia cultivars "Nightsky" == 0                 NA
## cultivarPetunia cultivars "Queen of Hearts" == 0          NA
## cultivarPhlox amplifolia == 0                             NA
## cultivarRudbeckia fulgida == 0                            NA
## cultivarSalvia farianacea == 0                            NA
## cultivarSalvia splendens "purple" == 0                    NA
## cultivarSalvia splendens "Red" == 0                       NA
## cultivarSalvia splendens "white" == 0                     NA
## cultivarSalvia yangii == 0                                NA
## cultivarSedum telephium == 0                              NA
## cultivarTagetes patula "orange" == 0                      NA
## cultivarTagetes patula "yellow" == 0                      NA
## (Adjusted p values reported -- BH method)
```

##### honey vs wild bees - pairwise comps

Going to use the raw count, because we're going to compare only within cultivar. Means we can use poisson distribution too

Going to only look at top 6 most attractive cultivars:

unique(data$cultivar)

```
# List of cultivars to filter
selected_cultivars <- c("Salvia yangii", "Aster ageratoides", "Eryngium planum \"Blauer Zwerg\"", "Echi

# Creating a subset dataset containing only the specified cultivars
top6cultivars <- bees_and_wildbees[bees_and_wildbees$cultivar %in% selected_cultivars, ]
```

make a figure, check all is in order

```
ggplot (top6cultivars , aes(x = cultivar, y = count, fill = insect_group))+
  scale_y_continuous(expand = c(0, 0)) + # forces X axis to 0, but in this case is overriden by ribbon
  coord_cartesian(ylim = c(0, 45)) +
  stat_summary(fun.y = "sum", geom = "bar", shape = 23, size = 3, alpha = 0.5) +
```

```
  theme_bw(12) +
  theme(axis.text.x = element_text(angle = 90, hjust = 1, vjust = 0.3)) +  # hjust and vjust nudge the 
  labs( x = "cultivar", y = "Total insects counted", title ="Sum count honey and wildbees, top 6 cutliva
```

```
## Warning: `fun.y` is deprecated. Use `fun` instead.

## Warning: Ignoring unknown parameters: shape
```

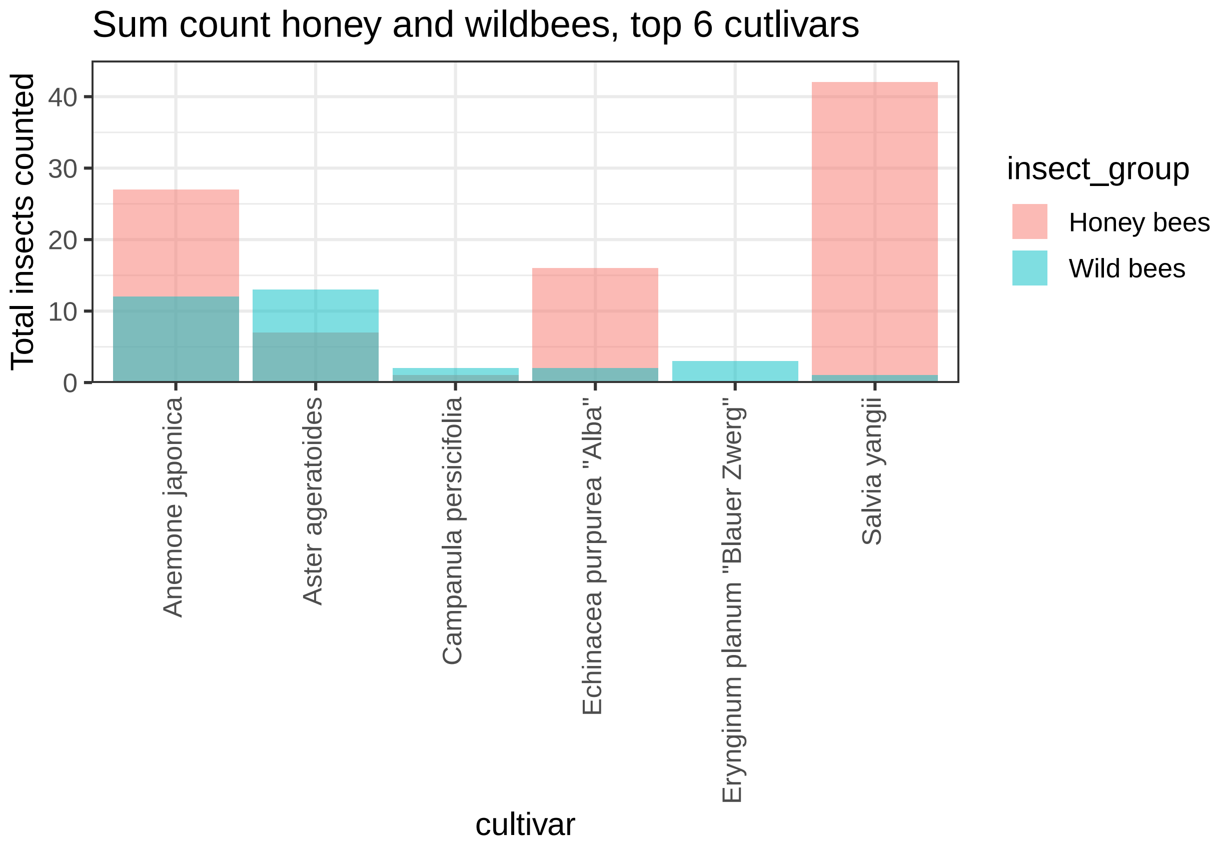

ok now lets see if we get do sensible modelling and pairwise comparisons with this

```
##NOTE IM USING THE RAW COUNT HERE, NOT THE M2 VARIABLE.

#model
m2beestop6 <- glmmTMB(count ~ as.factor(cultivar) * insect_group
             + (1 | location),
             ziformula=~1,
             family=poisson,
             data = top6cultivars)

Anova (m2beestop6)
```

```
## Analysis of Deviance Table (Type II Wald chisquare tests)
##
## Response: count
```

```
##                                      Chisq Df Pr(>Chisq)
## as.factor(cultivar)                 9.0092  5   0.108698
## insect_group                        6.8307  1   0.008960 **
## as.factor(cultivar):insect_group 19.5148  5   0.001541 **
## ---
## Signif. codes:  0 '***' 0.001 '**' 0.01 '*' 0.05 '.' 0.1 ' ' 1
```

```
#Dharma
simulateResiduals(m2beestop6, n = 500, plot = T) #dharma
```

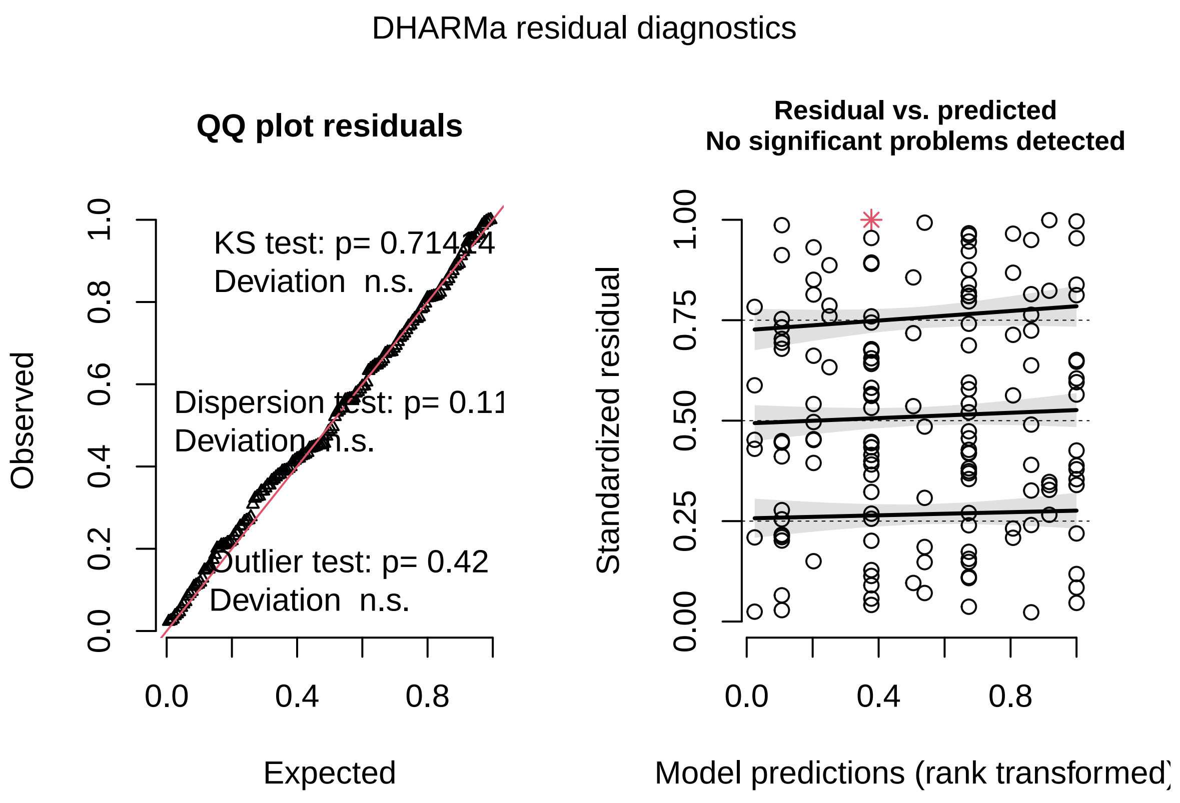

```
## Object of Class DHARMa with simulated residuals based on 500 simulations with refit = FALSE . See ?DI
##
## Scaled residual values: 0.7831745 0.9960342 0.3899917 0.8681415 0.3261663 0.8148226 0.3691393 0.42573
```

```
adjX <- glht(m2beestop6, alternative = "two.sided") # multcomp
summary(adjX, test = adjusted("BH"))
```

```
##
##   Simultaneous Tests for General Linear Hypotheses
##
## Fit: glmmTMB(formula = count ~ as.factor(cultivar) * insect_group +
##     (1 | location), data = top6cultivars, family = poisson, ziformula = ~1,
##     dispformula = ~1)
##
```

```
## Linear Hypotheses:
##                                                                       Estimate
## (Intercept) == 0                                                    -9.271e-02
## as.factor(cultivar)Aster ageratoides == 0                            8.551e-03
## as.factor(cultivar)Campanula persicifolia == 0                      -1.264e+00
## as.factor(cultivar)Echinacea purpurea "Alba" == 0                    5.316e-01
## as.factor(cultivar)Erynginum planum "Blauer Zwerg" == 0             -1.957e+01
## as.factor(cultivar)Salvia yangii == 0                                9.413e-01
## insect_groupWild bees == 0                                          -7.861e-01
## as.factor(cultivar)Aster ageratoides:insect_groupWild bees == 0      1.389e+00
## as.factor(cultivar)Campanula persicifolia:insect_groupWild bees == 0      1.451e+00
## as.factor(cultivar)Echinacea purpurea "Alba":insect_groupWild bees == 0    -1.266e+00
## as.factor(cultivar)Erynginum planum "Blauer Zwerg":insect_groupWild bees == 0  1.992e+01
## as.factor(cultivar)Salvia yangii:insect_groupWild bees == 0         -2.897e+00
##                                                                     Std. Error
## (Intercept) == 0                                                     2.922e-01
## as.factor(cultivar)Aster ageratoides == 0                            5.530e-01
## as.factor(cultivar)Campanula persicifolia == 0                       1.038e+00
## as.factor(cultivar)Echinacea purpurea "Alba" == 0                    3.520e-01
## as.factor(cultivar)Erynginum planum "Blauer Zwerg" == 0              8.003e+03
## as.factor(cultivar)Salvia yangii == 0                                3.252e-01
## insect_groupWild bees == 0                                           3.624e-01
## as.factor(cultivar)Aster ageratoides:insect_groupWild bees == 0      6.188e-01
## as.factor(cultivar)Campanula persicifolia:insect_groupWild bees == 0      1.300e+00
## as.factor(cultivar)Echinacea purpurea "Alba":insect_groupWild bees == 0     8.461e-01
## as.factor(cultivar)Erynginum planum "Blauer Zwerg":insect_groupWild bees == 0  8.003e+03
## as.factor(cultivar)Salvia yangii:insect_groupWild bees == 0          1.080e+00
##                                                                     z value
## (Intercept) == 0                                                     -0.317
## as.factor(cultivar)Aster ageratoides == 0                             0.015
## as.factor(cultivar)Campanula persicifolia == 0                       -1.217
## as.factor(cultivar)Echinacea purpurea "Alba" == 0                     1.510
## as.factor(cultivar)Erynginum planum "Blauer Zwerg" == 0              -0.002
## as.factor(cultivar)Salvia yangii == 0                                 2.894
## insect_groupWild bees == 0                                           -2.169
## as.factor(cultivar)Aster ageratoides:insect_groupWild bees == 0       2.244
## as.factor(cultivar)Campanula persicifolia:insect_groupWild bees == 0       1.116
## as.factor(cultivar)Echinacea purpurea "Alba":insect_groupWild bees == 0      -1.496
## as.factor(cultivar)Erynginum planum "Blauer Zwerg":insect_groupWild bees == 0   0.002
## as.factor(cultivar)Salvia yangii:insect_groupWild bees == 0          -2.682
##                                                                     Pr(>|z|)
## (Intercept) == 0                                                      0.9980
## as.factor(cultivar)Aster ageratoides == 0                             0.9980
## as.factor(cultivar)Campanula persicifolia == 0                        0.3831
## as.factor(cultivar)Echinacea purpurea "Alba" == 0                     0.2694
## as.factor(cultivar)Erynginum planum "Blauer Zwerg" == 0               0.9980
## as.factor(cultivar)Salvia yangii == 0                                 0.0439
## insect_groupWild bees == 0                                            0.0902
## as.factor(cultivar)Aster ageratoides:insect_groupWild bees == 0       0.0902
## as.factor(cultivar)Campanula persicifolia:insect_groupWild bees == 0       0.3964
## as.factor(cultivar)Echinacea purpurea "Alba":insect_groupWild bees == 0       0.2694
## as.factor(cultivar)Erynginum planum "Blauer Zwerg":insect_groupWild bees == 0   0.9980
## as.factor(cultivar)Salvia yangii:insect_groupWild bees == 0           0.0439
##
```

```
## (Intercept) == 0
## as.factor(cultivar)Aster ageratoides == 0
## as.factor(cultivar)Campanula persicifolia == 0
## as.factor(cultivar)Echinacea purpurea "Alba" == 0
## as.factor(cultivar)Eryngium planum "Blauer Zwerg" == 0
## as.factor(cultivar)Salvia yangii == 0                                   *
## insect_groupWild bees == 0                                              .
## as.factor(cultivar)Aster ageratoides:insect_groupWild bees == 0         .
## as.factor(cultivar)Campanula persicifolia:insect_groupWild bees == 0
## as.factor(cultivar)Echinacea purpurea "Alba":insect_groupWild bees == 0
## as.factor(cultivar)Eryngium planum "Blauer Zwerg":insect_groupWild bees == 0
## as.factor(cultivar)Salvia yangii:insect_groupWild bees == 0             *
## ---
## Signif. codes:  0 '***' 0.001 '**' 0.01 '*' 0.05 '.' 0.1 ' ' 1
## (Adjusted p values reported -- BH method)
```

```
#pairwise
meanie <- emmeans(m2beestop6, pairwise ~   insect_group|cultivar)
summary (meanie)
```

```
## $emmeans
## cultivar = Anemone japonica:
##  insect_group   emmean       SE  df  lower.CL  upper.CL
##  Honey bees    -0.0927    0.292 140 -6.70e-01     0.485
##  Wild bees     -0.8788    0.371 140 -1.61e+00    -0.146
##
## cultivar = Aster ageratoides:
##  insect_group   emmean       SE  df  lower.CL  upper.CL
##  Honey bees    -0.0842    0.551 140 -1.17e+00     1.004
##  Wild bees      0.5184    0.462 140 -3.95e-01     1.432
##
## cultivar = Campanula persicifolia:
##  insect_group   emmean       SE  df  lower.CL  upper.CL
##  Honey bees    -1.3566    1.045 140 -3.42e+00     0.710
##  Wild bees     -0.6912    0.760 140 -2.19e+00     0.812
##
## cultivar = Echinacea purpurea "Alba":
##  insect_group   emmean       SE  df  lower.CL  upper.CL
##  Honey bees     0.4389    0.342 140 -2.37e-01     1.115
##  Wild bees     -1.6128    0.758 140 -3.11e+00    -0.115
##
## cultivar = Eryngium planum "Blauer Zwerg":
##  insect_group   emmean       SE  df  lower.CL  upper.CL
##  Honey bees   -19.6665 8003.407 140 -1.58e+04 15803.500
##  Wild bees     -0.5284    0.699 140 -1.91e+00     0.853
##
## cultivar = Salvia yangii:
##  insect_group   emmean       SE  df  lower.CL  upper.CL
##  Honey bees     0.8486    0.269 140  3.18e-01     1.380
##  Wild bees     -2.8349    1.031 140 -4.87e+00    -0.796
##
## Results are given on the log (not the response) scale.
## Confidence level used: 0.95
##
```

```
## $contrasts
## cultivar = Anemone japonica:
##  contrast               estimate       SE  df t.ratio p.value
##  Honey bees - Wild bees    0.786    0.362 140   2.169  0.0318
##
## cultivar = Aster ageratoides:
##  contrast               estimate       SE  df t.ratio p.value
##  Honey bees - Wild bees   -0.603    0.499 140  -1.207  0.2296
##
## cultivar = Campanula persicifolia:
##  contrast               estimate       SE  df t.ratio p.value
##  Honey bees - Wild bees   -0.665    1.248 140  -0.533  0.5948
##
## cultivar = Echinacea purpurea "Alba":
##  contrast               estimate       SE  df t.ratio p.value
##  Honey bees - Wild bees    2.052    0.766 140   2.679  0.0083
##
## cultivar = Erynginum planum "Blauer Zwerg":
##  contrast               estimate       SE  df t.ratio p.value
##  Honey bees - Wild bees  -19.138 8003.407 140  -0.002  0.9981
##
## cultivar = Salvia yangii:
##  contrast               estimate       SE  df t.ratio p.value
##  Honey bees - Wild bees    3.684    1.020 140   3.612  0.0004
##
## Results are given on the log (not the response) scale.
```
