## Supplementary material for "High variability in the attractiveness of municipally-planted decorative plants to pollinators": table TS1

| Name in study | cultivars / plants observed | Annual / perennial / shurb |
| --- | --- | --- |
| Achillea filipendulina | Achillea filipendulina „Cloth of Gold“ | Perennial |
| Alcaltheae spp. | Alcaltheae x suffrutescens, Alcea rosea „Nigra“ | Biennial / Perrenial |
| Anemone japonica | Anemone japonica | Perennial |
| Aster ageratoides | Aster ageratoides "adustus Nanus" | Perennial |
| Begonia semperflorens "Eureka Scarlet" | Begonia semperflorens "Eureka Scarlet" | Perennial |
| Begonia semperflorens "Eureka White" | Begonia semperflorens "white" | Perennial |
| Bistorta affinis | Bistorta affinis | Perennial |
| Campanula persicifolia | Campanula persicifolia | Perennial |
| Canna indica | Canna indica | Perennial |
| Echinacea purpurea "pink" | Echinacea purpurea (unknown cultivars, probably 3 different cultivars) | Perennial |
| Echinacea purpurea „Alba“ | Echinacea purpurea „Alba“ | Perennial |
| Erigeron x cultorum | Erigeron x cultorum | Perennial |
| Erynginum planum „Blauer Zwerg“ | Erynginum planum „Blauer Zwerg“ | Perennial |
| Eryngium planum "Flachblatt-Mannstreu" | Eryngium planum "Flachblatt-Mannstreu" | Perennial |
| Galinsoga parviflora | Galinsoga parviflora | Perennial |
| Geranium x magnificum | Geranium x magnificum | Perennial |
| Hibiscus syriacus | Hibiscus syriacus | Perennial |
| Hydrangea macrophylla | Hydrangea macrophylla | Perennial |
| Nepeta grandiflora | Nepeta grandiflora | Perennial |
| Petunia cultivars "Nightsky" | Petunia cultivars "Nightsky" | Annual |
| Petunia cultivars "Queen of Hearts" | Petunia cultivars „Queen of Hearts“ | Annual |
| Phlox amplifolia | Phlox amplifolia „Minnehaha“ | Perennial |
| Rudbeckia fulgida | Rudbeckia fulgida „Little Gold Star“, Rudbeckia fulgida „Goldsturm“ | Perennial |
| Salvia splendens purple | Salvia splendens „Purple“ | Perennial |
| Salvia splendens red | Salvia splendens „Red“ | Perennial |
| Salvia splendens white | Salvia splendens „White“ | Perennial |
| Salvia yangii | Perovskia atriplicifolia „Blue Spire“ | Perennial |
| Sedum telephium | Sedum telephium | Perennial |
| Tagetes patula "orange" | Tagetes patula "Deep Orange" | Annual |
| Tagetes patula "yellow" | Tagetes patula „ Gold“, Tagetes patula „ Yellow“ | Annual |
